# KCC2 Enhancers Normalize Reflex Responses and Improve Locomotor Function after Chronic Spinal Cord Injury

**DOI:** 10.1101/2023.10.21.563363

**Authors:** Jadwiga N. Bilchak, Guillaume Caron, Simon M. Danner, Marie-Pascale Côté

## Abstract

Within a year after a spinal cord injury (SCI), 75% of individuals develop spasticity. While normal movement relies on the ability to adjust reflexes appropriately, and on reciprocal inhibition of antagonistic muscles, spastic individuals display hyperactive spinal reflexes and involuntary muscle co-contractions. Current anti-spastic medications can suppress uncontrolled movements, but by acting on GABAergic signalling, these medications lead to severe side-effects and weakened muscle force, making them incompatible with activity-based therapies. We have previously shown that pharmacologically enhancing activity of KCC2, a chloride cotransporter, reduces signs of spasticity in anesthetized chronic SCI rats. Here, we examine the effect of enhancing KCC2 in awake animals, using a battery of tests to assess multiple reflex pathways required for normal movement as well as locomotor function. Sprague-Dawley rats were implanted with chronic EMG electrodes bilaterally in ankle flexor and ankle extensor muscles and received a complete spinal transection at T12. Four weeks following SCI, the stretch reflex, the non-nociceptive cutaneous reflex pathway, the flexor withdrawal reflex, and the crossed-extensor reflex pathway as well as locomotor function were evaluated before and after receiving the KCC2 enhancer, CLP290. Our results show that enhancing KCC2 activity normalizes reflex responses in multiple pathways and reduces muscle co-contraction without weakening motor output, thereby improving stepping ability. This work reveals the substantial potential for KCC2 enhancers as a novel antispastic treatment.

## Introduction

Following a spinal cord injury (SCI), one of the major obstacles that individuals face is spasticity, a condition of disordered sensorimotor control (Nielsen et al., 2007). In addition to the pain and extreme discomfort it can induce, spasticity impairs residual motor function and hinders functional recovery. While activity-based therapies have beneficial effects on spasticity (Adam and Hicks, 2011; Elbasiouny et al., 2010; Field-Fote et al. 2022; Manella and Field-Fote 2013; Petropoulou et al., 2007), they are costly and often difficult or even impossible to implement for some patients, especially early after injury (Simon & Yelnik, 2010; Yelnik et al., 2009). Thus, it is necessary to use pharmacological treatments to mitigate spasticity. However, these medications often target GABAergic signalling, leading to severe systemic side effects. These include deep, long-lasting, and indiscriminate depression of spinal excitability as well as weakening of muscle force (Adams & Hicks, 2005; Lapeyre et al., 2010; Simon & Yelnik, 2010; Taricco et al., 2006; Thomas et al., 2010), both of which are detrimental to motor function and severely impede functional recovery. Rather than decreasing spasticity by depressing the overall excitability of spinal networks, a promising approach is to instead restore inhibition by targeting chloride homeostasis.

The chloride cotransporter, KCC2, has recently become recognized as a major player in the development of spasticity after SCI (Boulenguez et al., 2010). KCC2 is largely responsible for extruding chloride ions from neurons in the central nervous system and maintaining a high extracellular [Cl^-^] (Ben-Ari et al., 2012; Kaila et al., 2014; Payne et al., 2003). This allows for an influx of chloride ions in response to GABAergic and glycinergic signalling, and the post-synaptic inhibition that arises from this is vital for the function of multiple spinal pathways that regulate reflexes and locomotion. After SCI, KCC2 is progressively down-regulated in sublesional neurons, causing a depolarizing shift in the chloride reversal potential and a disruption of the hyperpolarization process (Bos et al., 2013; Boulenguez et al., 2010; Côté et al., 2014; Jean-Xavier et al., 2006; 2007; Hubner et al 2001). Interestingly, the reduction in spasticity observed with activity-based therapies relies on an increase in KCC2 expression (Chopek et al., 2015; Côté et al., 2014; Tashiro et al., 2015). This implicates KCC2 as a potential pharmacological target.

We have previously shown that the KCC2 enhancer, CLP257, mirrors the benefits of exercise and improves reflex modulation in terminal experiments conducted in chronic SCI rats (Bilchak et al., 2021a). In this study, by using chronically implanted EMG electrodes in awake animals, we examine the effects of pharmacologically enhancing KCC2 with CLP290 (a carbamate prodrug of CLP257 with a longer half-life) on multiple reflex pathways that are known to be disrupted with spasticity. Our results show that a single dose of CLP290 normalizes reflex responses towards those observed pre-injury, with differential effects on ankle flexor and ankle extensor muscles. We further tested whether CLP290 improves alternating locomotor patterns after chronic SCI. Our results reveal that enhancing KCC2 pharmacologically after SCI reduces co-contraction of antagonistic muscles, without dampening motor output, ultimately leading to improved locomotor function. Together, these results implicate enhancing KCC2 function as a promising strategy to mitigate spasticity while simultaneously improving locomotor recovery after SCI.

## Results

### Experimental design

In a rat model of complete thoracic SCI (T12), we investigated the effects of pharmacologically increasing KCC2 activity with CLP290 in awake, behaving animals. Rats (n=7) were bilaterally implanted with chronic electromyographic (EMG) electrodes in the tibialis anterior (TA), an ankle flexor, and the medial gastrocnemius (MG), an ankle extensor. EMG recordings were performed under the following conditions (**Figure 1**): 1) before injury (Pre-injury), 2) four weeks post-SCI before CLP administration (SCI - Pre CLP290), 3) five hours after receiving the KCC2 enhancer CLP290 (SCI - Post CLP290), and 4) after the effect of CLP290 was washed out, *i.e.,* 3 days later (SCI - Washout). We investigated the effect of CLP290 on four different spinal reflex pathways that are known to be disrupted in spasticity: the non-nociceptive cutaneous reflex pathway, the stretch reflex, the flexor withdrawal reflex, and the crossed extensor reflex pathway (**Figure 2**).

**Figure 1.**
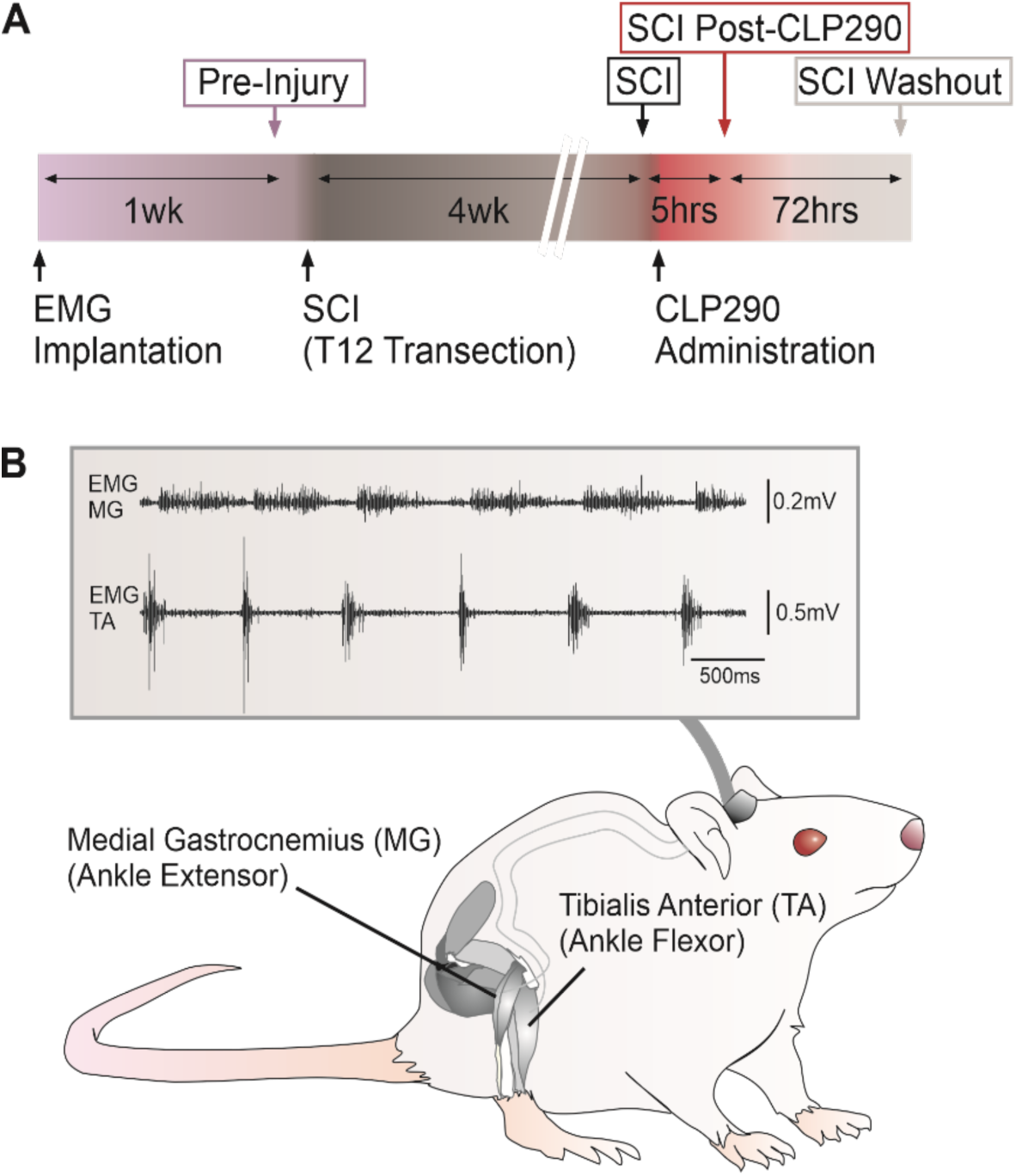
Experimental timeline and set-up. **A)** Timeline of EMG recordings (above) and surgical / pharmacological procedures (below). **B)** EMG electrodes were implanted bilaterally in the tibialis anterior (TA) and medial gastrocnemius (MG) muscles and EMG wires were led subcutaneously to a head connector.

**Figure 2.**
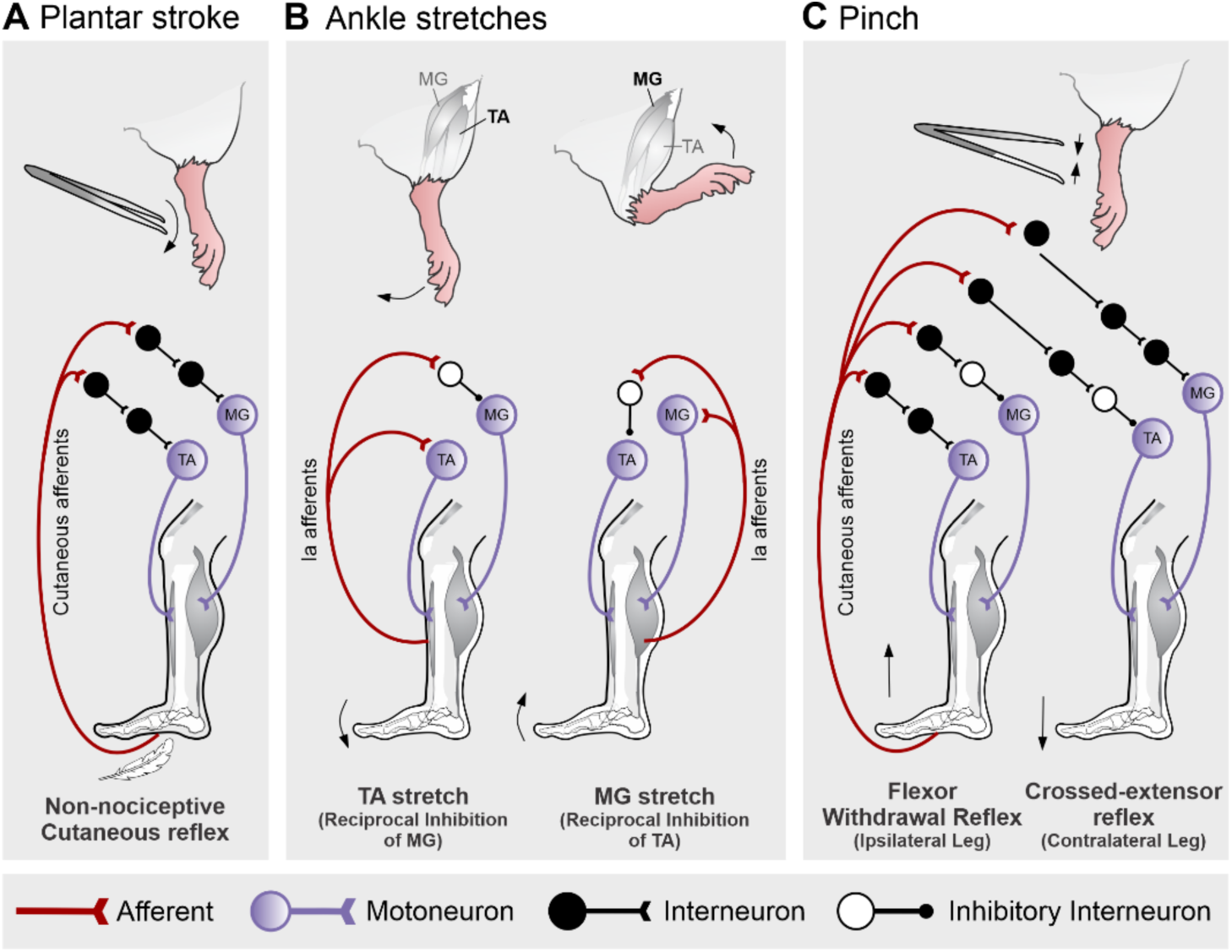
Methods to test reflex pathways. **A)** The cutaneous reflex pathway was activated by a gentle plantar stroke to activate non-nociceptive cutaneous afferents**. B)** The stretch reflex consisted of alternated plantarflexion followed by dorsiflexion to activate stretch receptors of TA and MG, respectively. **C)** The flexor withdrawal reflex and crossed-extensor reflex were activated by a mechanical noxious stimulus, *i.e.,* a brief pinch of the three medial toes along the proximal phalanges.

### Wind-up of Cutaneous Responses

The emergence of a reflex wind-up in response to repeated non-nociceptive stimuli has been described in both animal models (Garrison et al., 2011; Yates et al., 2011) and humans (Hornby et al., 2003; 2006) after SCI. This response is characterized by an increase in amplitude of EMG responses with each subsequent stimulus, suggesting an exaggeration of temporal facilitation, possibly due to injury-induced changes in motoneuronal intrinsic properties (Bennett et al., 2004). To investigate if CLP290 decreases exaggerated non-nociceptive cutaneous responses after SCI, a series of 15 gentle plantar strokes were performed to activate cutaneous afferents (**Figure 2A**) while the EMG response was recorded in MG and TA muscles. **Figure 3** depicts the EMG responses in MG and TA muscles in the same animal before injury, after chronic SCI, following CLP290 administration, and after washout. Before injury, consecutive strokes evoked responses of similar amplitude. After SCI, a reflex wind-up emerges with responses gradually increasing with each successive stimulus. CLP290 decreased the amplitude of the EMG responses and wind-up of cutaneous reflex. Three days later, once CLP290 had been washed out, the reflex wind-up returned. To characterize the modulation in response amplitude in MG and TA to the 15 consecutive strokes, we compared the slope of the wind-up between conditions for all animals. We found a significant difference in the slope for MG between conditions (*P* < 0.0001, **Figure 3C**). Consecutive strokes performed pre-injury produced responses of consistent amplitude in MG muscle with a slope close to 0 (0.0043 ± 0.005), similar to responses observed in healthy humans (Grey et al., 2008; Hornby et al., 2006). The slope increased following SCI (0.041 ± 0.011; *P* = 0.0089), suggesting the emergence of a reflex wind-up. CLP290 significantly reduced the slope (−0.0115 ± 0.009; *P* = 0.0009) as compared to SCI, resembling the slope observed pre-injury with values close to 0. The slope returned towards pre-CLP290 levels after washout and was significantly increased as compared to post-CLP290 (0.038 ± 0.009; *P* = 0.0002). This suggests that CLP290 decreases the cutaneous reflex wind-up observed after chronic SCI.

**Figure 3:**
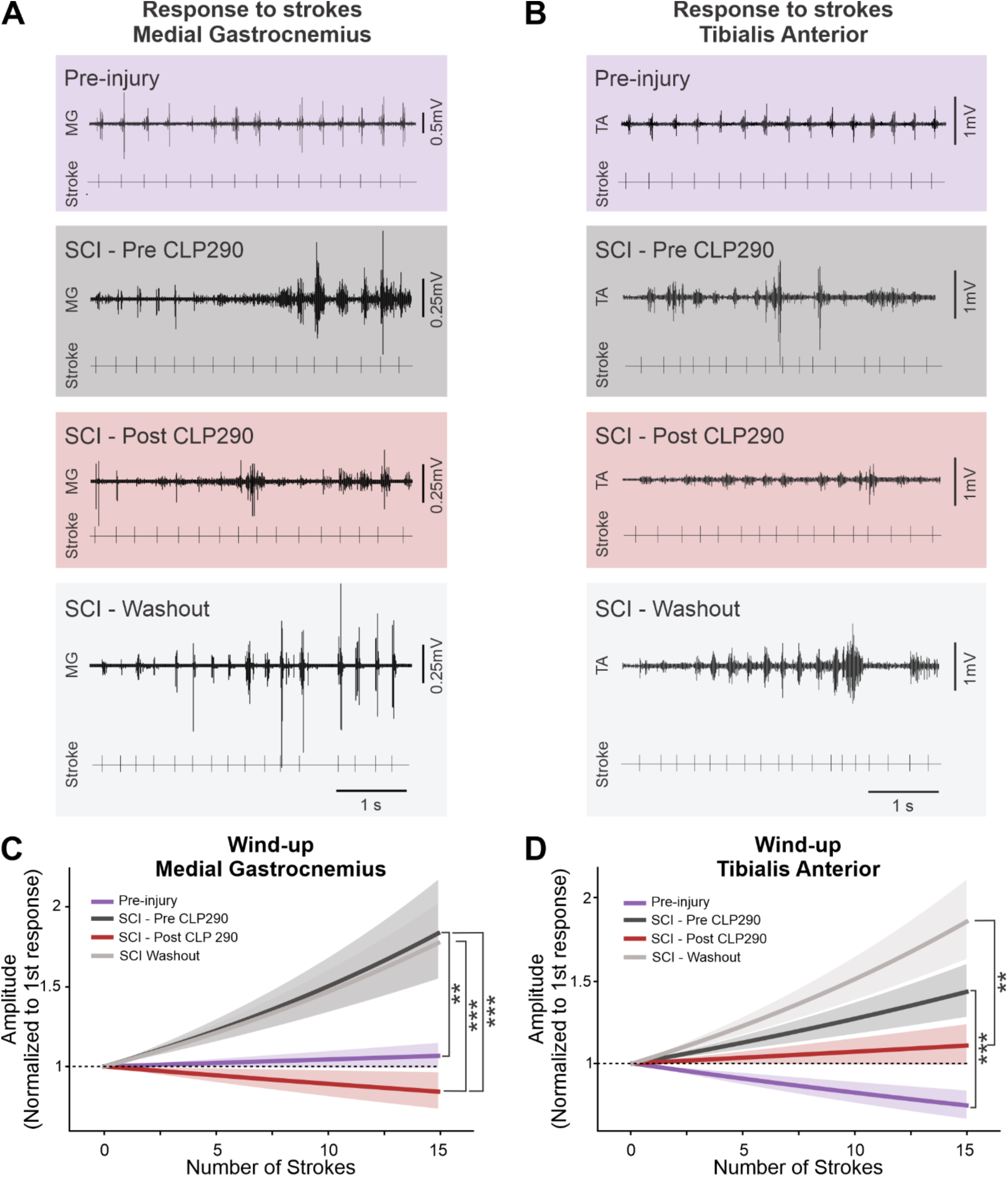
CLP290 reduces wind-up of cutaneous responses after SCI. **A-B)** Representative traces of consecutive cutaneous responses in MG (A) and TA (B) in the same animal before injury, 4 weeks post-SCI before CLP290, 4 weeks post-SCI after CLP290 administration, and after washout. **C)** After SCI, 15 consecutive strokes of the plantar surface induce a wind-up of the cutaneous reflex response, such that the slope of the response amplitude has a positive trend. After CLP290 is administered, the slope of responses no longer trends upward, resembling responses in animals before injury. **D)** The TA muscle shows similar wind-up responses as MG after chronic SCI, which is normalized by CLP290. Generalized mixed linear model. **p<0.01. ***p<0.001. (n = 6 rats, 12 TA muscles and 12 MG muscles).

Similarly, there was also a significant difference in slopes between conditions in TA (*P* < 0.0001). SCI increased the slope as compared to pre-injury (pre-injury: −0.0196 ± 0.008 vs SCI: 0.0241 ± 0.008; *P* = 0.0002, **Figure 3D**), suggesting the emergence of a reflex wind-up similar to that observed for MG. However, while there was a slight decrease in slope after administering CLP290, it did not reach statistical significance compared to SCI (0.0069 ± 0.007, *P* = 0.2502). Interestingly, the slope considerably increased after washout, beyond SCI values (0.0414 ± 0.009; *P* = 0.0069). This could indicate either a rebound effect following CLP290 treatment, or that reflex wind-up intensifies with time after SCI. Together, these results suggest that CLP290 reduces non-nociceptive reflex wind-up following SCI, with a particularly robust effect in ankle extensors.

### Stretches of Ankle Flexor and Extensor

Under normal circumstances, stretching of a muscle not only causes the reflex contraction of agonistic muscles, but also causes relaxation of antagonistic muscles. Following SCI, this reciprocal inhibition is diminished and can even be replaced by facilitation, resulting in muscle co-contraction (Boorman et al., 1996; Crone et al., 1994; Mirbagheri et al., 2014; Xia & Rymer, 2005). To investigate whether CLP290 decreases co-contractions after SCI, we performed EMG recordings in MG and TA in response to alternated stretches in plantarflexion and in dorsiflexion to activate stretch receptors (group Ia and II afferents) of TA and MG, respectively. A series of sequential stretches of the ankle were performed, with a stretch cycle consisting of a 0.25s stretch of the TA (plantarflexion) followed immediately by a 0.25s stretch of the MG (dorsiflexion) (**Figure 2B**). **Figure 4A** shows representative EMG traces of two stretch cycles from a single animal across all conditions. Prior to injury, TA and MG activity do not overlap, with TA activity occurring during the TA stretch and MG activity occurring during the MG stretch. To quantify the amount of co-contraction between MG and TA in all animals, we used the cosine similarity coefficient (**Figure 4B**). A cosine similarity of 0 indicates that the activity of TA and MG is alternated, whereas a cosine similarity of 1 indicates complete co-contraction. There was a significant difference in co-contraction across conditions (*P* = 0.0005), with an increase in co-contraction following SCI (pre-injury: 0.253 ± 0.154 vs. SCI: 0.496 ± 0.209; *P* = 0.0059). After administration of CLP290, MG-TA co-contraction values were close to pre-injury levels (0.332 ± 0.155) and significantly decreased compared to SCI (*P* = 0.0461). Co-contraction was returned to pre-CLP290 values after washout (0.463 ± 0.205) and significantly increased as compared to post-CLP290 (*P* = 0.0461). Together, this indicates that CLP290 reduces co-contraction between ankle flexors and extensors after SCI.

**Figure 4:**
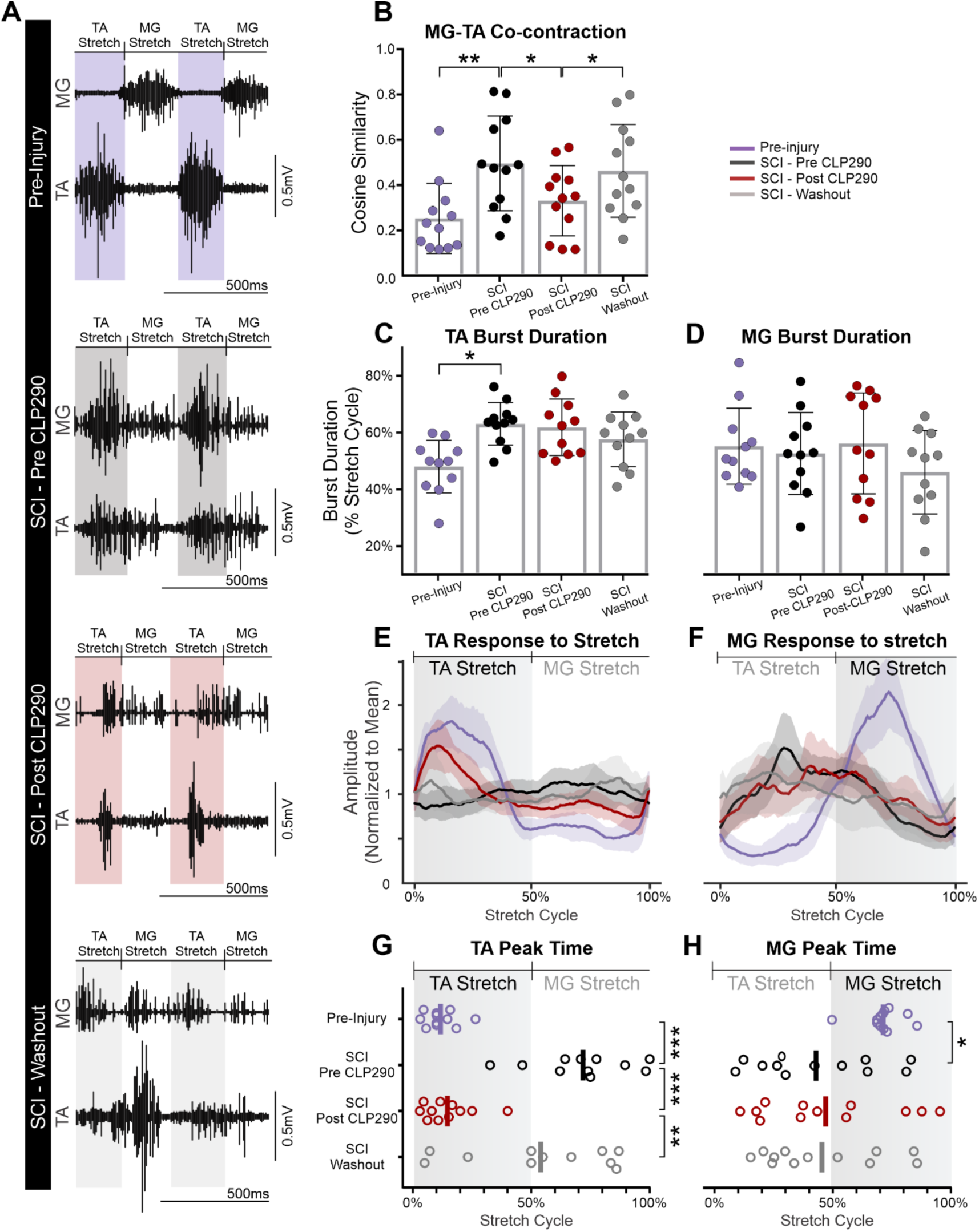
CLP290 reduces co-contraction of ankle flexor and extensor muscles during stretching. **A)** Representative traces of two stretch cycles from a single animal before injury, 4 weeks post-SCI before CLP290 administration, 4 weeks post-SCI after CLP290 administration, and after washout. **B)** The cosine-similarity of TA and MG activity significantly increases following SCI (p=0.005), indicating more co-contraction. Post CLP290, the cosine-similarity is significantly reduced (*P* = 0.034) back to pre-injury levels (n = 6 rats, 12 TA and 12 MG muscles) **C-D)** SCI increased TA (*P* = 0.01), but not MG burst duration, and CLP290 had no further effect on either TA nor MG (*P* =0.714 and *P* = 0.936, respectively) (n = 6 rats, 11 TA and 11 MG muscles) **E-H)** EMG envelopes for TA (E) and MG (F) averaged across all animals in response to stretches. Before SCI, peak TA activity occurs during the TA stretch (G), and the time of peak MG activity occurs during the MG stretch (H). TA (*P* < 0.001) and MG (p=0.02) peak times are significantly shifted. CLP290 shifted the peak response time towards pre-injury levels in TA (*P* < 0.001) but had no effect on MG (*P* = 0.126) (n = 6 rats, 11 TA and 12 MG muscles). B-D, Repeated-measure ANOVA; G, H, *Watson-Williams High Concentration F Test* *p<0.05, **p<0.01, ***p<0.001.

To explore the possible causes for this effect, we examined whether CLP290 altered burst duration in TA and MG. There was a significant interaction of burst duration and condition in TA (*P* = 0.0048) (**Figure 4C**). Although TA burst duration was increased following SCI (pre-injury: 48.1 ± 9.2% vs. SCI: 63.1 ± 7.5%, *P* = 0.01), CLP290 (61.9 ± 9.9%, *P* = 0.714) and washout had no further effect (57.7 ± 9.6%, *P* = 0.551). As for MG, no significant interaction between burst duration and condition was observed (*P* = 0.317) (**Figure 4D**). Together, this suggests that CLP290 does not affect TA or MG burst duration after injury.

Since the duration of MG and TA burst in response to stretch was not affected by CLP290, the increase in co-contraction could be explained by a shift in the timing of the response. To assess the timing of responses relative to TA stretches and MG stretches, we averaged all EMG responses for each animal and identified the relative timing of the stretch cycle at which the EMG response reached its maximum (**Figure 4E-F**). Before injury, TA activity reached its maximum during the TA stretch (12.3% of stretch cycle) (**Figure 4G**), and MG activity reached its maximum during the MG stretch (71.6% of stretch cycle) (**Figure 4H**). Peak timings were significantly shifted by SCI in both TA (63.1%, *P* = 0.0006) and MG (42.8%, *P* = 0.02). In TA, administration of CLP290 significantly shifted the response peak time back towards pre-injury levels (14.6%, *P* = 0.0002), an effect that was not present after washout (55.2%, *P* = 0.009). In contrast, the response peak time for MG was unaffected by CLP290 (46.9%, *P* = 0.126) and by washout (45.3%, *P* = 0.019). Overall, these results suggest that enhancing KCC2 activity with CLP290 reduces co-contraction of antagonistic ankle muscles, predominantly by shifting the timing of TA peak responses.

### Flexor Withdrawal Reflexes and Crossed-Extensor Reflexes

We then investigated the effect of CLP290 on nociceptive reflex pathways. To do so, we simultaneously recorded the flexor withdrawal reflex and the crossed-extensor reflex in response to a mechanical noxious stimulus, *i.e.,* a brief pinch of the three medial toes along the proximal phalanges. **Figure 5A** depicts the effect of CLP290 on these responses in the same animal. Before injury, the stimulus produced a robust response in the ipsilateral TA, while the ipsilateral MG remained silent (flexor withdrawal reflex). This was accompanied by a large response in the contralateral MG and silence in the contralateral TA (the crossed-extensor reflex). These reflexes allow the ipsilateral limb to withdraw from the noxious stimulus while the contralateral limb extends to maintain balance. After SCI, activity in both the ipsilateral MG and contralateral TA emerge, but to a much lesser extent after CLP290 administration. To quantify this, EMG amplitudes were compared within each muscle and across conditions. We first looked at the flexor withdrawal reflex, where the ipsilateral TA is active, and the ipsilateral MG is usually silent. There was no significant interaction between EMG amplitude and conditions in the ipsilateral TA (*P* = 0.175, **Figure 5B**) indicating that EMG amplitude in these muscles was unaffected by SCI and/or by CLP290. However, there was a significant interaction between EMG amplitude and conditions in the ipsilateral MG (*P* = 0.011, **Figure 5C**). The ipsilateral MG amplitude increased following SCI (3.25 ± 2.187, *P* = 0.0199). However, while there was a modest decrease in amplitude following CLP290, the effect was not significant (2.448 ± 1.355, *P* = 0.210) and the amplitude remained unchanged after washout (3.977 ± 3.265, *P* = 0.170). We then looked at the crossed-extensor reflex, where the contralateral MG is active, and the contralateral TA is usually silent. There was no significant interaction of EMG amplitude and conditions in the contralateral MG (*P* = 0.320, **Figure 5D**) indicating that EMG amplitude in these muscles was unaffected by SCI and by CLP290. However, there was a significant interaction in the contralateral TA (*P* = 0.0004, **Figure 5E).** The amplitude of the contralateral TA was increased after SCI (2.60 ± 1.41) as compared to pre-Injury (*P* = 0.0037). More importantly, CLP290 decreased its amplitude as compared to SCI (1.37 ± 0.908, *P* = 0.0022). After washout, the amplitude increased again (*P* = 0.0009), returning to pre-CLP290 values (2.39 ± 1.286). Together, this suggests that SCI disrupts both the flexor withdrawal reflex and crossed-extensor reflex due to a lack of inhibition on the ipsilateral MG and the contralateral TA. In addition, enhancing KCC2 activity with CLP290 restored inhibition in the contralateral TA, but had little effect on the ipsilateral MG.

**Figure 5:**
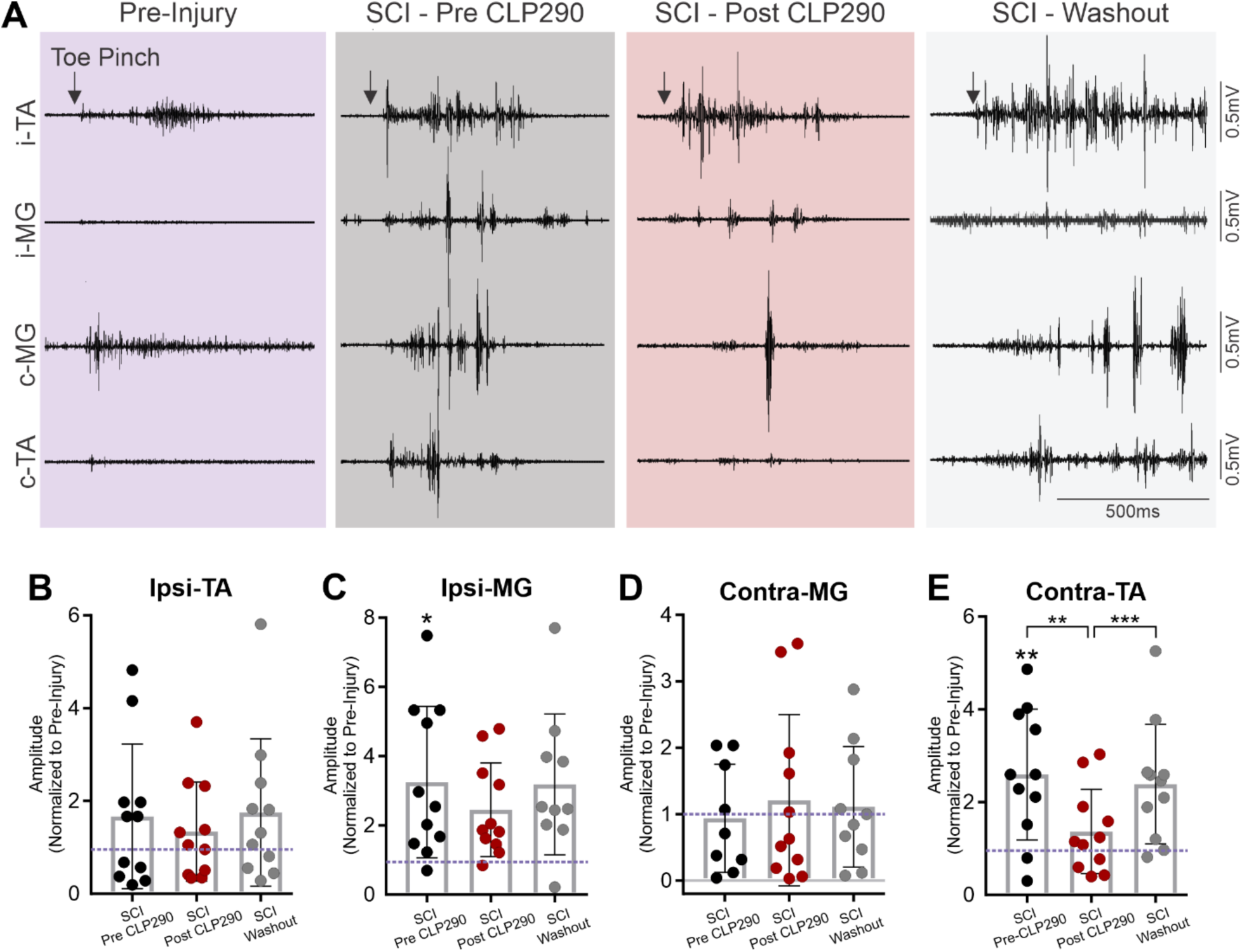
CLP290 restores flexor inhibition in the crossed-extensor reflex. **A)** Representative traces of pinch responses in the same animal before injury, 4 weeks post-injury before and after CLP290 administration, and after washout. **B)** The ipsilateral TA shows no significant change in response amplitude across conditions; **C)** Similarly, there is no significant effect of SCI or CLP290 application on the response amplitudes in the ipsilateral MG. **D)** A loss of inhibition is observed in the contralateral MG, with SCI significantly increasing response amplitude (p = 0.019). This, however, was not affected by CLP290 treatment (p = 0.21) **E)** SCI also significantly increases response amplitude in the contralateral TA (p = 0.0037), suggesting a loss of inhibition acting on the contralateral TA motoneurons. CLP290 restored this inhibition and significantly reduced response amplitude towards pre-injury levels (p = 0.0022). (n=6 rats, 11 TA and 11 MG muscles). Repeated measure ANOVA. *p<0.05. **p<0.01. ***p<0.001

### Intralimb and Interlimb Coordination during Treadmill Stepping

We further assessed how CLP290 affects intra- and inter-limb coordination during locomotion after SCI. **Figure 6A** shows representative EMG traces from a single animal during treadmill stepping across all conditions. Intralimb (MG vs TA) and interlimb (MG vs MG, or TA vs TA) co-contraction was estimated using the cosine similarity coefficient. For intralimb coordination, there was a significant interaction between cosine similarity and conditions (P < 0.0001). SCI increased the cosine similarity, *i.e.,* co-contractions between MG and TA within the same limb (pre-injury: 0.228 ± 0.09 vs. SCI: 0.419 ± 0.160; *P* = 0.0043, **Figure 6B**). CLP290 significantly reduced intralimb co-contraction (0.275 ± 0.12, *P* = 0.0043), and co-contraction resumed after washout (0.467 ± 0.17, *P* = 0.0043). Assessing similarity between TA muscles in the two limbs, there was a trending increase in co-contraction after SCI which appeared to normalize with administration of CLP290 (**Figure 6C**). However, the differences failed to reach significance across groups (*P* = 0.1407). Co-contraction between MG muscles also showed no significant interaction and indicated no changes following SCI or CLP290 administration (*P* = 0.4707, **Figure 6D**). Together, these results suggest that enhancing KCC2 function restores the intralimb muscle alternation that was lost after SCI and may also have a small effect on interlimb coordination between flexor muscles.

**Figure 6:**
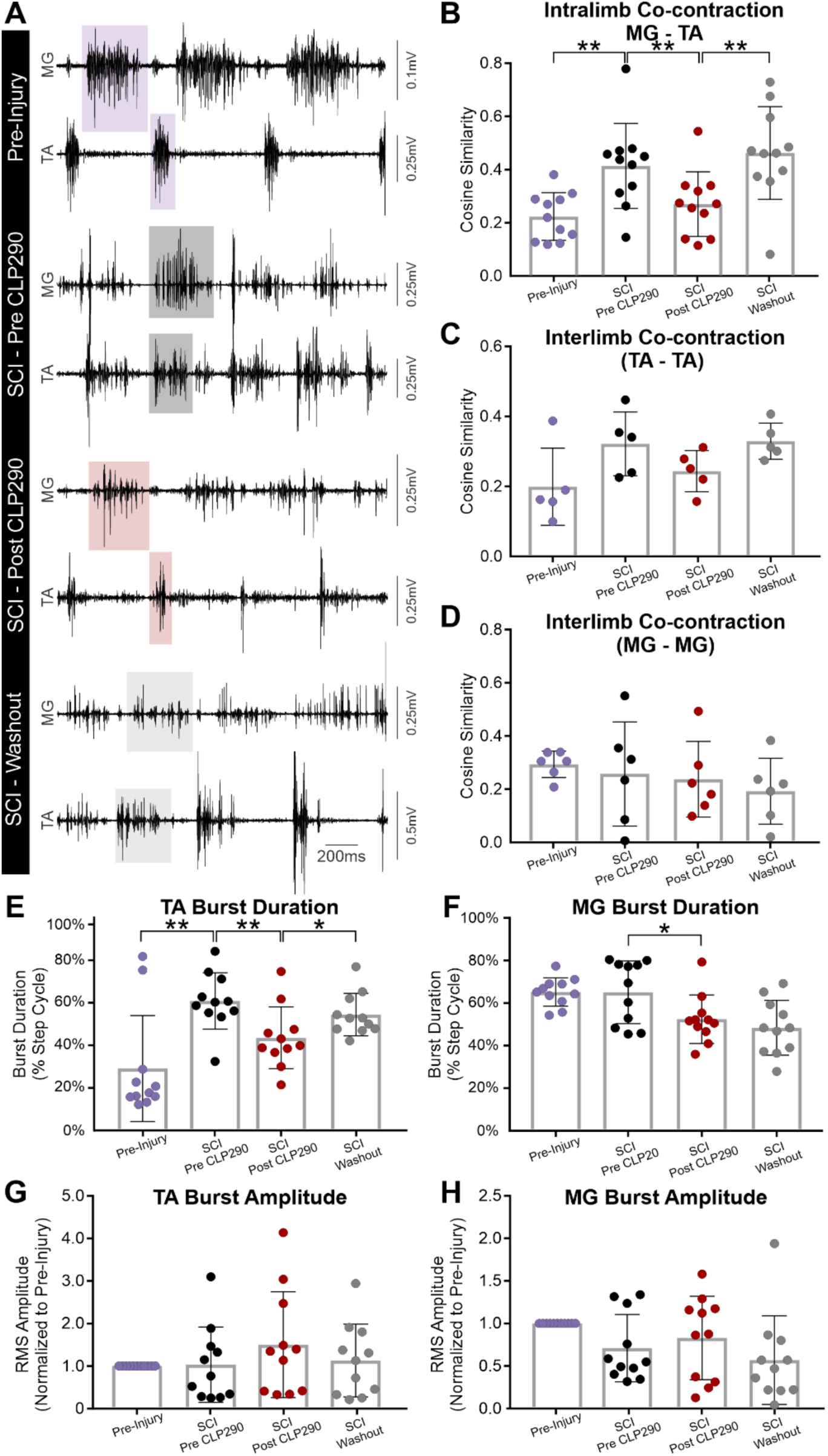
CLP290 improves locomotor function. **A)** Representative traces of treadmill stepping from a single animal before injury, 4 weeks post injury before CLP290 and after CLP290 administration, and after washout. **B)** After SCI, there is a significant increase in the cosine similarity between TA and MG within each leg (p=0.0043, n=6 rats, 11 TA and 11 MG muscles) indicating increased co-contraction of these antagonistic muscles. Administration of CLP290 reduced the cosine similarity back towards pre-injury levels (p=0.0043). **C-D)** There were no significant changes in cosine similarity between the left and right TA muscles (C) nor the left and right MG muscles (D) (n=6 rats, 5 pairs of TA muscles and 6 pairs of MG muscles) **E)** TA burst duration was significantly longer following SCI (p=0.0072) and was reduced towards pre-injury levels by CLP290 (p=0.0038) (n=6 rats, 11 TA muscles) **F)** In contrast, the average MG burst duration remained unchanged by SCI, but was modestly reduced by CLP290 (n=6 rats, 11 MG muscles). **G-H)** Importantly, the changes observed in locomotion were not accompanied by significant changes in burst amplitude in either the TA (G) nor the MG (H) (n=6 rats, 11 TA and 11 MG muscles). Repeated measure ANOVA. *p<0.05. **p<0.01.

We next examined the effect of SCI and CLP290 on burst duration. There was a significant interaction between burst duration and conditions for TA (*P* = 0.0029, **Figure 6E**). TA burst duration increased after SCI (pre-Injury: 29.2 ± 25.0% vs. SCI: 60.8 ± 13.3%, *P* = 0.0072), while CLP significantly reduced this increase (43.5 ± 14.5%, *P* = 0.0038). Following Washout, burst duration returned towards SCI levels (54.5 ± 10.0%) and significantly increased as compared to after CLP290 administration (*P* = 0.0214). There was also a significant interaction between burst duration and conditions for MG (*P* = 0.0010, **Figure 6F**), although MG burst duration was not significantly changed following SCI (pre-Injury: 65.2 ± 6.7% vs SCI: 65.1 ± 14.7%, *P* = 0.975). MG burst duration was reduced by CLP290 (52.4 ± 11.4%, *P* = 0.027) but remained unchanged following Washout (48.4 ± 12.9%, p = 0.027). Together, this suggests that the effect of CLP290 on co-contraction during locomotion may be due to a normalization of TA burst duration, along with a slight decrease in MG burst duration.

Finally, EMG amplitude, normalized to pre-injury values, was assessed to determine how CLP290 affects motor output during locomotion. There was no significant interaction between EMG amplitude and condition in either TA (*P* = 0.3158, **Figure 6G**) or MG (*P* = 0.0637, **Figure 6H**). This indicates that the improvements in locomotor pattern observed with CLP290 are not accompanied by the dampening of motor output seen with Baclofen and other GABAergic agonists currently used to treat spasticity (Angeli et al., 2012; Dario and Tomei, 2004; Dietz et al., 2022; Elovic, 2001; Kirshblum, 1999).

## Discussion

Cation chloride cotransporters have recently been implicated in the disruption of synaptic inhibition in multiple pathologies, including developmental disorders such as epilepsy (Sullivan and Kadam, 2020), autism (Ferrari and Ben-Ari Y, 2020), and Down syndrome (Savardi et al., 2020) and psychiatric disorders such as schizophrenia, depression, and stress (Maguire et al., 2020). In the spinal cord, substantial work has been conducted in the dorsal horn on understanding the contribution of chloride homeostasis to neuropathic pain (Kahle et al., 2014). Enhancing KCC2 function has proven very successful in this context, with CLP290 administration normalizing mechanical withdrawal thresholds in models of peripheral nerve injury (Gagnon et al., 2013) and morphine-induced hyperalgesia (Ferrini et al., 2017). Moreover, unlike equipotent doses of other analgesics, CLP290 showed no effect on motor performance in the rotarod assay (Gagnon et al., 2013). This provided evidence that modulating Cl^-^ homeostasis does not depress motor function and, thus, may be a viable method to treat spasticity after SCI without impeding locomotor recovery. This was further corroborated in a staggered hemisection model of SCI in which CLP290 re-established the excitatory-inhibitory ratio of pathways residing between the two lesions, thus improving overground stepping by facilitating the relay of brain-derived commands (Chen et al., 2018).

Here, we examined multiple spinal pathways involved in the control of movement and how restoring Cl^-^ homeostasis affects reflex modulation and locomotor function after a complete SCI. Our results show that increasing KCC2 activity after SCI normalizes reflex responses towards pre-injury levels and, by decreasing co-contraction of muscles without diminishing motor output, improves locomotor function. In addition, we observed differential effects in ankle flexors and extensors following injury and administration of CLP290, providing an unexpected glimpse into how SCI affects chloride-mediated inhibition in these distinct motoneuronal populations.

### Enhancing KCC2 Activity Reduces Wind-up of Cutaneous Responses

The wind-up of reflex responses is a well-known phenomenon that occurs after SCI and is attributed to an increase in temporal summation (Hornby et al., 2003; 2006). This likely stems from the reappearance of persistent inward currents (PICs) in the chronic stages of injury. PICs are voltage gated Ca^2+^ and Na^+^ currents that amplify and prolong responses to brief descending commands, allowing spinal motoneurons to produce sustained voluntary muscle contractions (Brownstone et al., 1994; Heckmann et al., 2005). In the acute stages of spinal injury, PICs are absent due to the abrupt lack of descending modulation but are slowly revived as the injured cord adapts to its new-found isolation from supraspinal centers (Button et al., 2008; Li et al., 2004; Murray et al., 2010). Once this occurs, the PICs are both more easily triggered (Baker & Chandler, 1987; Dietz & Sinkjaer, 2007) and more difficult to terminate (Bennett et al., 2004), resulting in long-lasting, debilitating muscle spasms (Bennett et al., 2004; Gorassini et al., 2004; Li et al., 2004). In our experiments, consecutive cutaneous stimuli produced responses of increasing amplitude following SCI. After increasing KCC2 activity with CLP290, response amplitudes remained consistent throughout, resembling those observed pre-injury. This could have resulted from a restoration of chloride-mediated hyperpolarization in motoneurons, thus allowing PICs to be terminated more easily by inhibitory inputs.

The fact that the MG muscle responded more robustly to CLP290 treatment further suggests that CLP290 has a differential effect on flexor and extensor motoneurons, as is the case with activity-based therapies (Chopek et al., 2015). However, the wind-up response involves a complex set of pathways subject to state-dependent modulation, and while an inability to suppress PICs is a possible contributor, other mechanisms are likely at play.

### Enhancing KCC2 Activity Reduces Co-contraction of Flexor and Extensor Muscles During Stretching

While it is considered likely that reciprocal inhibition is impaired after SCI and is a significant contributor to the pathophysiology of spasticity (Boorman et al., 1996; Elbasiouny et al., 2010; Morita et al., 2001; Nielsen et al., 2007; Xia and Rymer, 2005), its precise cause after SCI remains unclear. The disruption of descending input, collateral sprouting, and changes in presynaptic inhibition are all likely involved (Crone et al., 2003; Elbasiouny et al., 2010). There is further evidence suggesting that changes in chloride homeostasis could also play a significant role. During development, the shift from reciprocal excitation to reciprocal inhibition coincides closely with the rise of chloride-mediated inhibition (Delpy et al., 2008; Wang et al., 2008). Moreover, in an *in-vitro* preparation, that shift towards reciprocal inhibition can be reversed back toward excitation by down-regulating or pharmacologically inhibiting KCC2 (Gackiere & Vinay, 2015). Our results show that pharmacologically enhancing KCC2 activity reduces co-contraction of ankle flexor and extensor muscles after SCI, providing evidence that impaired chloride homeostasis contributes to the injury-induced disruption of reciprocal inhibition. While there was no change in burst duration following CLP290 treatment, the burst activity in the TA was consolidated and shifted such that the majority of activity took place during the TA stretch, as it did pre-injury. The remaining tonic component that contributed to the long burst duration, however, appeared to be unaffected by CLP290. In the MG, CLP290 produced no shift in response timing, further suggesting a disparity in how KCC2 influences inhibition in flexor and extensor motoneurons. Together, this suggests that enhancing KCC2 activity restores reciprocal inhibition by acting on TA motoneurons after chronic SCI.

### Enhancing KCC2 Activity Restores Flexor Inhibition in the Crossed-Extensor Reflex

Excess excitability of the flexor withdrawal reflex has been shown to arise concurrently with increased muscle tone and spasms after acute SCI in humans (Hiersemenzel et al., 2000). But at later stages, once clinical signs of spasticity have developed, the amplitude of the flexor withdrawal reflex decreases (Hiersemenzel et al., 2000; Knikou & Conway, 2005; Muller & Dietz, 2006), and the resultant leg-joint torques are smaller than in healthy individuals (Deutsch et al., 2005). It was, thus, postulated that activity of the flexor withdrawal reflex is unrelated to the pathophysiology of spasticity (Dietz & Sinkjaer, 2007). Here, we show that there is no significant change in TA response to an ipsilateral pinch in the chronic stages of injury, but rather an increase in excitation of the ipsilateral MG, which would normally be silenced by inhibitory interneurons. This suggests that the perceived decrease in flexor withdrawal reflex excitability may alternatively be attributed to a lack of inhibition acting on MG motoneurons. Surprisingly, CLP290 had little to no effect on this, implying the involvement of another mechanism unrelated to KCC2 expression. Potential causes could be the lack of serotonergic modulation after injury (Aggelopoulos et al., 1996) and/ or an injury-induced decrease in presynaptic inhibition of MG afferents, which would unlikely be affected by CLP290 since primary afferents do not express KCC2 (Kanaka et al., 2001; Mao et al., 2012).

We also observed a marked increase in activity of the contralateral TA following SCI, similar to the crossed excitation seen in spinal cats (Frigon et al., 2012). In contrast to the ipsilateral MG response, this activity was significantly reduced by CLP290 towards pre-injury levels. This suggests that the crossed excitation that arises after injury is due in part to an impairment of chloride-mediated inhibition in flexor motoneurons and that acute treatment with a KCC2 enhancer can restore the crossed-extensor reflex. Given that the interneurons involved in the flexor and crossed-extensor reflex are postulated to be a part of the central pattern generator for locomotion (Guertin, 2012; Jankowska et al., 1967a, 1967b), restoring normal inhibition in these pathways could lead to improvements in locomotor recovery.

### Enhancing KCC2 Activity Improves Locomotor Function

The alternation of muscle activity during locomotion relies on alternating periods of excitatory and inhibitory inputs (Pratt & Jordan, 1987; Shefchyk & Jordan, 1985). For this alternation to take place between extensors and flexors and between both hindlimbs, there must be mutual inhibition, which is heavily reliant on GABA (Cazalets et al., 1994; Cowley & Schmidt, 1995; Grillner & Jessell, 2009; Kremer & Lev-Tov, 1998). Given the major influence chloride homeostasis has on GABAergic signalling, a long-lasting down-regulation of KCC2 after SCI is likely to lead to abnormal motoneuronal responses to inhibitory inputs (Stil et al., 2011), which would ultimately impair locomotion. While CLP290 was shown to improve locomotion in a staggered hemisection mouse model of SCI mostly via an increase in interneuronal relay transmission between the two lesions (Chen et al., 2018), our results suggest a larger role for restoration of motoneuronal chloride homeostasis when there is no remaining supraspinal inputs. Increasing KCC2 activity restored alternation between the MG and TA muscles within each limb and reinstated inhibition to control the duration of muscle bursting, particularly in the TA muscle. Interestingly, these results (i.e., decreased co-contraction and burst duration) resemble those seen after successful body-weight supported step-training in both animal models (Barbeau & Rossignol, 1991) and in humans (Gorassini et al., 2009), suggesting that pharmacologically manipulating chloride homeostasis may reproduce some of the improvements associated with exercise-based rehabilitation (Bilchak et al., 2021a-b; Caron et al., 2023). Our results further suggest that by acting to restore endogenous inhibition solely in the central nervous system (Medina et al., 2014; Rivera et al., 1999), CLP290 may circumvent the muscle weakness and depression of motor output seen with current anti-spastic medications as burst amplitude during stepping remained unaffected by CLP290 administration. Together, this indicates that CLP290 improves reciprocal inhibition between TA and MG motoneurons within each limb, resulting in improved alternation between the antagonistic muscles without a decrease in motor output.

### KCC2 Enhancers Differentially Affect Ankle Flexor and Extensor Muscles

Flexor and extensor motoneurons respond differently both to spinal transection and to exercise-based rehabilitation (Chopek et al., 2014; 2015; Skup et al., 2012). It is, therefore, likely that treatment with CLP290 would affect KCC2 membrane expression in the two motoneuronal populations differently. Since CLP290 only affects pre-existing KCC2 by facilitating its insertion in the membrane (Gagnon et al., 2013), CLP290 could potentially have a lesser effect on a set of neurons that suffer a substantial decrease in KCC2 expression after injury. Alternatively, a particular neuronal population may already have sufficient KCC2 membrane expression, in which case, CLP290 is unlikely to produce functional differences, as demonstrated in SCI animals that already displayed increased KCC2 membrane expression due to exercise (Bilchak et al., 2021a). A previous study examining KCC2 mRNA levels after transection SCI found that extensor motoneurons largely retain KCC2 expression, while flexor motoneurons show the more substantial decrease (Chopek et al., 2015). However, since both the MG and the TA responded to CLP290 treatment in at least one of our experiments, it seems unlikely that their differences rely solely on the efficacy of CLP290 or on KCC2 expression. Instead, the differences in efficacy appear to be task-dependent, suggesting that flexors and extensors rely on chloride-mediated inhibition for different tasks after SCI. Where CLP290 had no effect, the changes in muscle activity following SCI could depend on other mechanisms such as dendritic sprouting or presynaptic inhibition of primary afferents, which is highly task- and state-dependent (Côté & Gossard, 2003) and impervious to KCC2 activity (Kanaka et al., 2001; Mao et al., 2012).

### Conclusions

We have shown here that restoring chloride homeostasis after SCI positively affects multiple reflex modalities in the awake animal, namely cutaneous, noxious, and stretch responses, offering insights into the critical role of KCC2 activity on sublesional spinal networks after SCI. With this holistic approach, we aimed to capture the multifaceted nature of spasticity (McKay et al., 2018; Pandyan et al., 2005) while avoiding the effects of anaesthesia for more clinical relevance.

One of the most notable observations is that a single dose of CLP290 increases alternation between antagonistic muscles during stepping, resulting in improved locomotion. These encouraging results implicate CLP290 as a potential anti-spastic medication that 1) could be provided acutely after injury, contrary to activity-based therapies and 2) does not exert a dampening effect on motor output as other anti-spastic medications do. Given that CLP290 increases inhibition only in certain pathways, a combinatorial approach with activity-based therapies may provide significant advantages in the chronic phase of SCI, when the implementation of a rehabilitation program or stimulation-based therapies is feasible.

## Materials and Methods

### Experimental Design

In a rat model of complete thoracic SCI (T12), we investigated the effects of pharmacologically increasing KCC2 activity on spasticity and locomotor ability in awake, behaving animals. Rats (n=7) were implanted with chronic electromyograph (EMG) electrodes bilaterally in the tibialis anterior (TA) and medial gastrocnemius (MG). EMG responses to stretch, cutaneous and nociceptive stimuli, as well as stepping ability on a treadmill, were evaluated in the intact animal (pre-Injury) before a complete spinal transection at T12 spinal level was performed. On the fourth week following injury, spasticity and locomotor recovery were evaluated before (SCI - Pre CLP290) and 5h after receiving an intraperitoneal injection of the KCC2 enhancer, CLP290 (SCI Post-CLP290). Washout recordings were conducted 72h after drug administration (SCI - Washout). All procedures were performed in accordance with protocols approved by Drexel University College of Medicine Institutional Animal Care and Use Committee (IACUC). All animal work followed National Institutes of Health guidelines for the care and use of laboratory animals and complied with Animal Research: Reporting of In Vivo Experiments (ARRIVE).

### Surgical procedures and postoperative care

Adult female Sprague Dawley rats (240-300g, Charles River Laboratories) were housed in pairs and kept in a 12-h light-dark cycle with controlled room temperature and ad libitum access to food. On the day of EMG electrode implantation, rats were anesthetized with isoflurane (1-4%) in O_2,_ their hindlimbs and head were shaved, and disinfected, and small incisions were made over the skull, tibialis anterior (TA, ankle flexor) muscles, and medial gastrocnemius (MG, ankle extensor) muscles (**Figure 1B**). EMG wires (AS632, Cooner Wire) were then drawn subcutaneously from the head incision to each of the leg incisions. Prior to surgery, 3mm of insulation was removed from the end of each electrode wire, and wire pairs were sutured together such that the two bared regions were 5mm apart. Each pair of electrode wires was inserted into the belly of its respective muscle with a curved needle and sutured in place. To ensure electrodes were correctly placed and operative, an isolated pulse stimulator (A-M Systems) delivered single bipolar pulses (100 µsec) to each wire pair while hindlimb muscles were observed for activity. Correct electrode placement was also verified post-mortem. The ground electrode was placed subcutaneously along the back. Following electrode insertion, three stainless steel screws were inserted into the skull and the electrode head connector was fixed to the screws using dental cement. Once the connector was secured, all skin incisions were cleansed and sutured together, fastened with wound clips, and treated with New Skin. Animals received a single injection of slow-release buprenorphine (0.05 mg/kg, s.c.), and were removed from anaesthesia. Following electrode implantation, rats were singly housed and received daily saline (3ml, s.c.) and Baytril (15mg/kg, s.c.) for 3 days.

One week after EMG implantation, rats underwent a complete spinal transection at the low thoracic level (T12) as described previously (Beverungen et al., 2020; Bilchak et al., 2021a; Caron et al., 2020; Côté et al., 2014). Briefly, rats were anesthetized with isoflurane (1-4%) in O_2_ and, under aseptic conditions, a laminectomy was performed at the T10–T11 vertebral level. The dura was carefully slit open, the spinal cord completely severed with small scissors, and the cavity filled with absorbable hemostats (Pfizer, New York, NY, USA) to promote hemostasis. The completeness of the lesion was ensured by the distinctive retraction of the rostral and caudal spinal tissue and by examining the ventral floor of the spinal canal during surgery and was confirmed post-mortem. Paravertebral muscles were sutured, and the skin closed with wound clips. Upon completion of the surgery, animals received a single injection of slow-release buprenorphine (0.05 mg/kg, s.c.), and daily saline (5ml, s.c.) and Baytril (15mg/kg, s.c.) for 7 days to prevent dehydration and infection, respectively. Bladders were expressed manually at least twice daily until the voiding reflex returned.

### Drug Preparation

Less than an hour before administration, the KCC2 enhancer CLP290 (generous gift from Dr. Y. de Koninck, Université Laval, Qc, Canada) was suspended in 40% hydroxypropyl-beta-cyclodextrin (HPCD) to make a 20mg/ml suspension. This was sonicated in a sonicating bath for ten minutes, then diluted 1:1 in deionized H_2_O to create a 10mg/ml solution. The final dose of CLP290 was 100mg/kg. Following another ten minutes of sonication, the solution was drawn into a syringe and delivered to rats i.p. immediately after their “Pre-CLP290” recording and five hours before their “Post-CLP290” recording. As the half-life of CLP290 is approximately 5 hours (Ferrini et al., 2017; Gagnon et al., 2013), the 5-hour gap allowed time for CLP290 to take full effect while also allowing rats to rest between recording sessions.

### EMG recordings

Five days after EMG implantation, animals were acclimated to the treadmill for 20 minutes while running the treadmill at various speeds. Animals were also familiarized with each of the sensorimotor response assays. EMG responses were recorded in rats one day before injury (Pre-injury), 4 weeks after injury, prior to receiving CLP290 (SCI), 5 hours after receiving CLP290 (SCI post-CLP290), and 72 hours after receiving CLP290 (SCI Washout). Throughout all conditions, the same experimenter performed the assays and handled rats during recordings. Apart from Pre-Injury, the experimenter was blind to which condition (SCI, SCI Post-CLP290, or SCI Washout) was being recorded. Rats were briefly anesthetized with isoflurane (1-4%) in O_2_ to attach the tether onto the rat’s head connector. EMG activity obtained during the experiment was amplified (100-1000×; A-M Systems, Carlsborg, WA), band-pass filtered (10–5000 Hz) and the signal was digitized (10 kHz) before being stored on a computer for analysis in Spike2 version 8 (Cambridge Electronic Design Limited, Cambridge, UK). Once it was verified that EMG signal was stable in all channels, anaesthesia was removed. Experimenters waited at least 10 minutes before beginning the recording to ensure rats were calm and had regained full responsiveness.

Four assays were performed: cutaneous response wind-up, alternating stretches, nociceptive responses, and stepping ability. For cutaneous responses, rats were held firmly on their side while both hindlimbs rested so that muscles were fully unloaded. Once the animal was calm and hindlimb EMG activity was silent, a blunted forceps of 0.5 mm width was used to stroke along the plantar surface of the paw, from the heel to the toes. The applied pressure was set to not move the paw or ankle. On each leg, a series of fifteen consecutive strokes was performed at an approximate frequency of 4 Hz. Alternating stretches were performed by holding rats firmly by their chest, with their hindlimbs hanging freely, until they were calm and EMG activity was silent. By pressing gently on the plantar surface of one paw, a maximal dorsiflexion of the foot was produced for 0.25 s, thus stretching the MG muscle. This was immediately followed by a 0.25 s long maximal plantar flexion, which stretched the TA muscle (Cabaj et al., 2017; Slawinska & Kasicki, 2002). The two stretches, each lasting 0.25 s, together formed a 0.5 s long stretch cycle. Ten consecutive stretch cycles were performed in each leg. For nociceptive responses, rats were held firmly on their side while both hindlimbs rested so that muscles were fully unloaded. Curved forceps of 0.5mm width were used to pinch the three medial toes along the proximal phalanges. Bipedal stepping was performed on a motorized treadmill at 15 cm/s, with perineal stimulation and manual assistance provided to support the hindquarters and maintain lateral stability. Statistical analysis was conducted in Python, R, and in GraphPad Prism. Analyses and statistical tests used for individual measurements are described within each section.

### Cutaneous Response Wind-up Analysis

The response to each of the 15 strokes was defined within a 0.25 s time window starting at stroke onset, and the root-mean squared (RMS) values for each response were measured. To assess response wind-up, we used a generalized linear mixed model with a gamma distribution on log-transformed data for each muscle (TA and MG). The glmmTMB package (Brooks et al., 2017) for R 3.6 was used to fit the model. Error distributions and link-functions were chosen to satisfy model assumptions, as confirmed by quantile–quantile (Q–Q) plots and histograms of Pearson’s residuals. A fixed effect for conditions, a covariate for stroke number, and their interaction were included. The model controlled for interindividual differences, using a per animal and hindlimb random intercept. Random slopes were included for conditions with an unstructured covariance structure and for stroke number with a first order auto-regressive covariant structure. A full factorial dispersion model was also included. *Post-hoc* comparisons using Tukey’s test were used to compare differences of the slope of the covariate stroke number between conditions.

### Alternating Stretch Analysis

Following recordings, EMG signals were rectified and smoothed with a second-order Savitzky–Golay filter and a fixed window of 100ms. Stretch cycle onsets were defined as the beginning of TA stretch. The maximum EMG amplitude within each stretch cycle was defined as the peak amplitude within the cycle minus the minimum amplitude within the cycle. Burst durations were defined as the percentage of the stretch cycle in which an EMG burst was >20% of the maximum EMG amplitude. Signal less than 0.01 mV in amplitude was excluded as noise. Burst durations were calculated for each muscle and averaged across the 10 stretch cycles performed in each hindlimb. The averaged values were compared across conditions using a one-way repeated measures ANOVA, followed by a Holm-Sidak *post-hoc* test. Unless otherwise stated, values were normally distributed as assessed by Shapiro-Wilk’s test (p > 0.05) and Q-Q plot. There was homogeneity of variances, as assessed by Levene’s test of homogeneity of variance (p > 0.05). Mauchly’s test of sphericity indicated that the assumption of sphericity was met (p > 0.05). Where it was violated, a Greenhouse-Geisser correction was applied. Outliers were identified and excluded using the ROUT method with Q set to 1% (Motulsky & Brown, 2006).

Co-contraction between the MG and the TA muscles in each hindlimb was defined by the cosine similarity of their filtered EMG signals. The cosine similarity function compares the orientation of two vectors (i.e., the two filtered EMG signals) and provides an estimation of similarity that is independent of amplitude, without the need to define burst durations. The similarity values between the MG signal and TA signal in each hindlimb were then compared across conditions using a repeated measures ANOVA followed by a Holm-Sidak *post-hoc* test.

To assess the timing of TA and MG responses within stretch cycles, EMG responses from all 10 cycles performed in each hindlimb were averaged to create an average response envelope. The time-point in the stretch cycle at which the averaged response envelope reached its maximal amplitude was defined as the “peak response time”. Due to the cyclical nature of the stretch cycles, the Watson-Williams High Concentration F-Test was used to compare differences in peak response times across groups.

### Nociceptive Response Analysis

EMG activity was rectified and smoothed with the Savitzky–Golay filter as stated above, and the RMS amplitude of each muscle’s response was measured within a 1.5 second time-window following pinch onset (defined as the moment that the forceps made contact with the toes). Four pinches per leg (two legs per rat) were analyzed and their responses averaged. To account for large differences in EMG amplitude between different muscles and different animals, response amplitudes were normalized to pre-injury values. A repeated measures one-way ANOVA was used to compare the normalized amplitudes across conditions, followed by Holm-Sidak *post-hoc* comparisons.

### Locomotor Function Analysis

EMG activity from twenty consecutive steps was extracted, rectified, and smoothed with a Savitzky–Golay filter as stated above. The beginning of each step cycle was defined as the onset of the flexion response. Burst durations for each muscle were defined in the same manner as for the alternating stretch analysis and averaged across all 20 step-cycles. The averaged values were compared across conditions using a repeated measures ANOVA followed by a Holm-Sidak *post-hoc* test. Co-contraction was calculated between the MG and the TA muscles in each hindlimb, between the left and right TA muscles, and between the left and right MG muscles. This was defined by the cosine similarity of their filtered EMG signals, as conducted for the analysis of alternating stretches. The similarity values were then compared across conditions using a repeated measures ANOVA followed by a Holm-Sidak *post-hoc* test. Burst amplitudes for each muscle were defined as the RMS amplitude of filtered and rectified EMG responses within each step-cycle. These were averaged across all 20 step-cycles for each condition, normalized to pre-injury values, then compared across conditions using a repeated measures ANOVA followed by Holm-Sidak *post-hoc* test.

## Acknowledgments

We thank Drs. Yves de Koninck and Annie Castonguay from Université Laval for the generous gift of CLP290 and technical assistance. This work was supported by grants from the National Institute of Neurological Disorders and Stroke (R01 NS083666) and the Craig H. Neilsen Foundation (189758).

